# The smooth-walled human RVOT contains trabeculations that cause conduction delay

**DOI:** 10.1101/2023.11.04.565628

**Authors:** Bjarke Jensen, Fernanda M. Bosada, Michiel Blok, Koen T Scholman, Igor R Efimov, Bastiaan J Boukens

## Abstract

**Aims:** The right ventricular outflow tract (RVOT) is the outlet from the right ventricle and is the initiating substrate of life-threatening arrhythmias. While the luminal wall of the RVOT is often assumed to be without the complex trabecular meshwork that characterizes the right ventricular free wall, the anatomy of the RVOT is an understudied subject. Our aim was to investigate whether trabeculations occur in the RVOT and to assess whether this impacts electrical propagation.

**Methods & Results:** We used high-resolution MRI and serial sectioning to reconstruct the macroscopic details of the human RVOT and identified cases exhibiting much trabeculation. The smooth lumen of the RVOT varied between 9% and 23% of the total RV anterior surface (N=11). Histological analysis on additional six hearts indicated that the RVOT compact layer is thinner when trabeculations are present. RNA sequencing of four human donor hearts revealed enrichment in the subendocardial region of 88 genes associated with cardiac conduction and trabeculations (P adjusted<0.05). Finally, we selected two human donor hearts showing trabeculations in the RVOT from which we generated wedge preparation and performed optical and electrical mapping. The trabecular regions demonstrated high degree of fractionation when compared to non-trabeculated regions, which coincided with delayed activation.

**Conclusion:** Trabeculations are found in the RVOT, and their extent varies among individuals. This impacts on the thickness of the compact wall in the RVOT, restricting the depth of tissue at which clinical interventions can be performed, as well as influencing electrical propagation and possible arrhythmogenicity.

## Introduction

The right ventricular outflow tract (RVOT) is the outlet from the right ventricle of the heart and it has received much attention since it was discovered in early 2000s that it is the origin of life-threatening arrhythmias in patients with Brugada syndrome (1, 2). The reason why these lethal arrhythmias originate in the myocardial wall of the RVOT has been the subject of research ever since. Genetic screening and experimental studies have led to the hypothesis that there may be a developmental basis for these arrhythmias, which would explain the hereditary nature of Brugada syndrome (3-6). Whether the presence of cardiac conduction tissue in the RVOT or potential variation in RVOT myocardium size between individuals contribute to arrhythmias initiation is unclear.

The luminal side of the left and right ventricular free wall is trabeculated, whereas the luminal side of the outflow tract wall of both ventricles is considered void of trabeculations and is smooth. Much remains to be elucidated on how these regional differences in trabeculations develop. In fact, the processes that influence the degree of trabeculation may not be entirely similar between the right and left ventricles (7), particularly at the level of the outflow tracts (8). We have previously posited that the degree of incorporation of the smooth embryonic outflow tract myocardium may be a driver of functional and morphological variation in the RVOT (9). However, little is known about the morphological variation of the RVOT. At least in the pig, the extent of RVOT trabeculation varies between hearts (10) and among mammals the RVOT varies in length and size (11).

The objective of this study was to examine the diversity of RVOT morphology in adult human hearts and to assess the functional implications thereof. Our morphological descriptions are complemented by bulk RNA sequencing and electrical mapping. In summary, the present findings reveal an unrecognized variation in RVOT trabeculation and show how this results in a propensity for conduction delay especially at higher pacing frequencies.

## Materials and Methods

### Ethical statement

The hearts used for histological characterization were provided by the archive of the Department of Medical Biology, VU University Medical Centre, Amsterdam. As the nature of the histological characterization was retrospective and the hearts were anonymized, we did not require informed consent as described in our institutional ethical guidelines and the tenets of the Declaration of Helsinki. The donor hearts used for RNA sequencing, electrophysiological characterization and 3D reconstruction were provided by Mid-America Transplant Services (Saint Louis, MO). The procedures in this study were in accordance with the legal requirements for the use of donor hearts in the USA. The ethics committee (IRB) of the Washington University School of Medicine approved the use of non-transplantable de-identified hearts for research purposes.

### Collecting hearts from the Mid-America Transplant Service St Louis, USA

Explanted human hearts obtained from Mid-America Transplant Services (Saint Louis, MO, USA) from deceased donors whose hearts were deemed unsuitable for transplantation. The arrested heart was maintained at 4-7° C to preserve the tissue during the 15-20 minute transportation from the operating room to the research laboratory. All explanted hearts were cardioplegically arrested in the operating room following cross-clamping of the aorta. For each heart, a total of 100mls of cardioplegic solution (in mmol/L: NaCl 110, CaCl2 1.2, KCl 16, MgCl_2_ 16, NaHCO_3_ 10; 4°C) was flushed through each of the right and left coronary arteries in the operating room immediately after receiving the heart from the surgical team.

### 3-D reconstruction of human heart using MRI

From a previous study (7), we used the MRI-generated image stack of an *ex vivo* normal heart from a 51 year-old male who died of pancreatic adenocarcinoma. The spatial resolution was 0.5 × 0.5 × 0.5 mm^3^ and thus substantially better than current clinical resolution (12). Briefly, the image stack was imported to the 3D software Amira (version 3D 2021.2, FEI SAS, Thermo Fisher Scientific). Labeling of various structures including the RVOT were done in the Segmentation Editor module. From the label file, using the Generate Surface function, a surface file was made that was visualized using the Surface View function. The conversion of the 3D model to an interactive pdf was done as previously described (13).

### 3-D reconstruction of tissue block based on bright field photographs

A single heart was used from which we collected the RVOT. The tissue was cryo-preserved and send for section to BioInVision, Inc, Cleveland Ohio (www.bioninvision.nl). The CryoViz™ cryo-imaging system consists of a digital cryo-microtome housed within a − 20 °C freezer chamber, a microscopic imaging system consisting of a high NA objective, a low-noise, cooled camera, a robotic XYZ positioner, and Programmable Logic Controller (PLC)-based control. The section thickness was 40 μm and section pick-up 10 μm. The image stack was imported to the 3D software Amira (version 3D 2021.2, FEI SAS, Thermo Fisher Scientific). Labeling of various structures including the RVOT were done in the Segmentation Editor module. From the label file, using the Generate Surface function, a surface file was made that was visualized using the Surface View function.

### Determining variation in size of smooth-walled RVOT

To quantify the variation in the size of the smooth part of the RVOT, we made screenshots of lumen casts of 11 hearts from the Atlas of Human Cardiac Anatomy from University of Minnesota (Specimen code, sex, and age are listed in Supplementary Table 1). All measurements were done in ImageJ 1.51k (NIH, USA) using the ‘Polygon selections’ tool for areas and the ‘Segmented Line’ tool for lengths. On the screenshots of the lumen casts of the 11 hearts from the Atlas of Human Cardiac Anatomy, we measured on the RV lumen cast the area which had a smooth surface (RVOT) and the total RV area. The RVOT area was then calculated as a percentage of total RV area per heart. On the histology samples, we measured the average width of both the compact and trabecular wall.

### Gross morphology and Histology

From our archives, we used hearts that had been fixed in formalin and later stored in preservation fluid I (aqua, glycerin, ethanol, methanol and phenol, Orphi Farma, NL). Two hearts had been set aside during a study of ventricular trabeculations (7), as having a normally trabeculated RVOT (T92-11007.3) and as having a much trabeculated RVOT (T92-9196.3). These hearts were photographed macroscopically. We selected six hearts of people who died of causes not related to the heart and isolated transmural blocks of chamber wall for histology and morphometric analyses (Sex, age, and cause of death are listed in Supplementary Table 2). These hearts were initially intact, and we were therefore blind to the luminal anatomy of the RVOT. To guide the isolation of blocks, we first inserted a probe from the pulmonary artery to the apical part of the right ventricular cavity. The approximately 2cm long block that was isolated along the probe and contained the pulmonary valve is referred to as the RVOT. The RV free wall sample was isolated halfway between the RV apex and the pulmonary valve. All isolated blocks were imbedded in paraplast. Microtome cut sections of 10 μm thickness were mounted on glass and stained with Masson’s trichriome (muscle is red, collagen is blue). Images were taken at a resolution of 0.97 μm/pixel and using the Photomerge tool (Layout, Auto) in Adobe Photoshop (version 13.0.1, Adobe Systems Incorporated, USA) we generated stitched images of 30252x7665 pixels.

### Morphometric analysis of the RVOT sections

The average width was calculated by dividing the area of the wall (compact wall only, or compact and trabeculated wall together) with the length of the compact wall. The length was measured as the midline (parallel to the epicardium) through the compact wall. This midline was later used to divide the epicardial and endocardial halves of the compact wall and measure myocardial density in the two halves. Myocardial density was measured as average grey scale value for epicardial and endocardial halves on images that had been converted to 8-bit (max signal value 255) and inverted such that lumen, interstitium, and fat would give 0 or values very close to 0. Walls of coronary vasculature yielded values well above 0, but these tissues were a tiny fraction of all tissues. The base of the pulmonary artery was surrounded by a turret, or cuff, of myocardium. We defined the width of the base of the turret as the distance from the lowest part of the pulmonary arterial valve hinge to the epicardial-most part of the ventricular wall. The height of the turret was then the distance from the base of the turret to the superior-most attachment of myocardium to the wall of the pulmonary artery.

### RNA isolation and sequencing

Four hearts were used from which we collected transmural samples of the RVOT. Total RNA isolation was performed on approximately 100 mg myocardium. In short, the tissue was homogenized using an ultra-Thurrax homogenizer. Whole tissue RNA was isolated using the Trizol method, followed by a DNase treatment and purification using RNeasy MinElute kit (Qiagen). RNA quality, concentration, and RNA integrity number (RIN) scores were assessed on the 2100 Bioanalyzer (Agilent Technologies). Samples with RIN scores above 6.8 were selected for RNA-seq. cDNA libraries were amplified using TruSeq RNA library prep kit v2 (Illumina) in accordance with manufacturer’s instructions using 50 ng / 0.5 ug of RNA as input. Sequencing was performed on Illumina HiSeq4000 instrument (50 bp single reads). Reads were aligned to hg19 built of the human transcriptome with STAR (14). Differential expression analysis was performed with edgeR (15). PANTHER (16) was used for gene onthology (GO) biological process analysis. Benjamini-Hochberg correction was performed for multiple testing-controlled p values and statistically significant enriched terms were functionally grouped and visualized.

### Optical mapping of RVOT preparation

Two hearts were used for optical and electrical characterization. We generated a wedge preparation from the right ventricle, including the RVOT, by removing the atria, left ventricle and posterior right ventricular wall. We cannulated both the right and left coronary artery. Following the dissection, the preparations were placed into the optical mapping setup and perfused at 37°C with Tyrode’s solution (in mmol/L) 128.2 NaCl, 4.7 KCl, 1.19 Na H2PO4, 1.05 MgCl2, 1.3 CaCl2, 27.0 NaHCO3, and 11.1 glucose. pH was maintained at 7.4 by equilibration with a mixture of 95% O2 and 5% CO2. For recording of optical action potentials, we stained the preparations by perfusing with 10 μM di-4-ANEPPS (Molecular Probes, Eugene, OR) for 10 min. The excitation-contraction uncoupler blebbistatin (10-15 μM, Tocris Bioscience, Ellisville, MO) was added to the perfusate to remove motion artifacts. Excitation light was delivered by a 520±5 nm light emitting diode (Prizmatix, Southfield, MI) and emitted fluorescence was filtered > 610 nm and recorded by a CMOS-sensor, (100 x 100 elements, 1 kHz, MICAM Ultima, SciMedia Ltd, Costa Mesa CA). Local bipolar electrograms were recorded at various location and stimulation frequencies (PowerLab 26T; AD-Instruments, Colorado Springs, CO).

## Results

### Anatomical evidence for trabeculation of the free wall of the RVOT

Figure 1 shows a MRI-based 3-dimensional reconstruction of a normal human heart. The right ventricular wall was trabeculated on the luminal side (Figure 1A), whereas the anterior superior part was smooth showing the RVOT. Unexpectedly, the anterior-right aspect of the RVOT luminal side was marked by trabeculations which extended across the entire length of the wall (Figure 1C). Macroscopic examination of one of the hearts revealed almost complete trabeculation of the RVOT myocardial wall (Figure 2A-B). The trabeculations reached the base of the right-facing and the non-facing leaflets of the pulmonary valve and the anterior portion of the RVOT that was smooth-walled was ±2 mm. In the other 5 hearts, the anterior smooth-walled portion was ±2 cm long. To determine the variation in the anterior smooth-walled surface of RVOT myocardium in more detail, we analyzed the lumen casts of 11 hearts from the Atlas of Human Cardiac Anatomy. In these hearts the RVOT accounted for ±14% of the right ventricular area from the anterior view while considerable variation existing between individual hearts (9-23%) (Figure 2C-D). In a previous study from our group, the degree of trabeculation of the right and left ventricle were not correlated (7) and a very substantial smooth-walled RVOT could be found even in hearts with an extremely high degree of LV trabeculations (Figure 2E).

**Figure 1.**
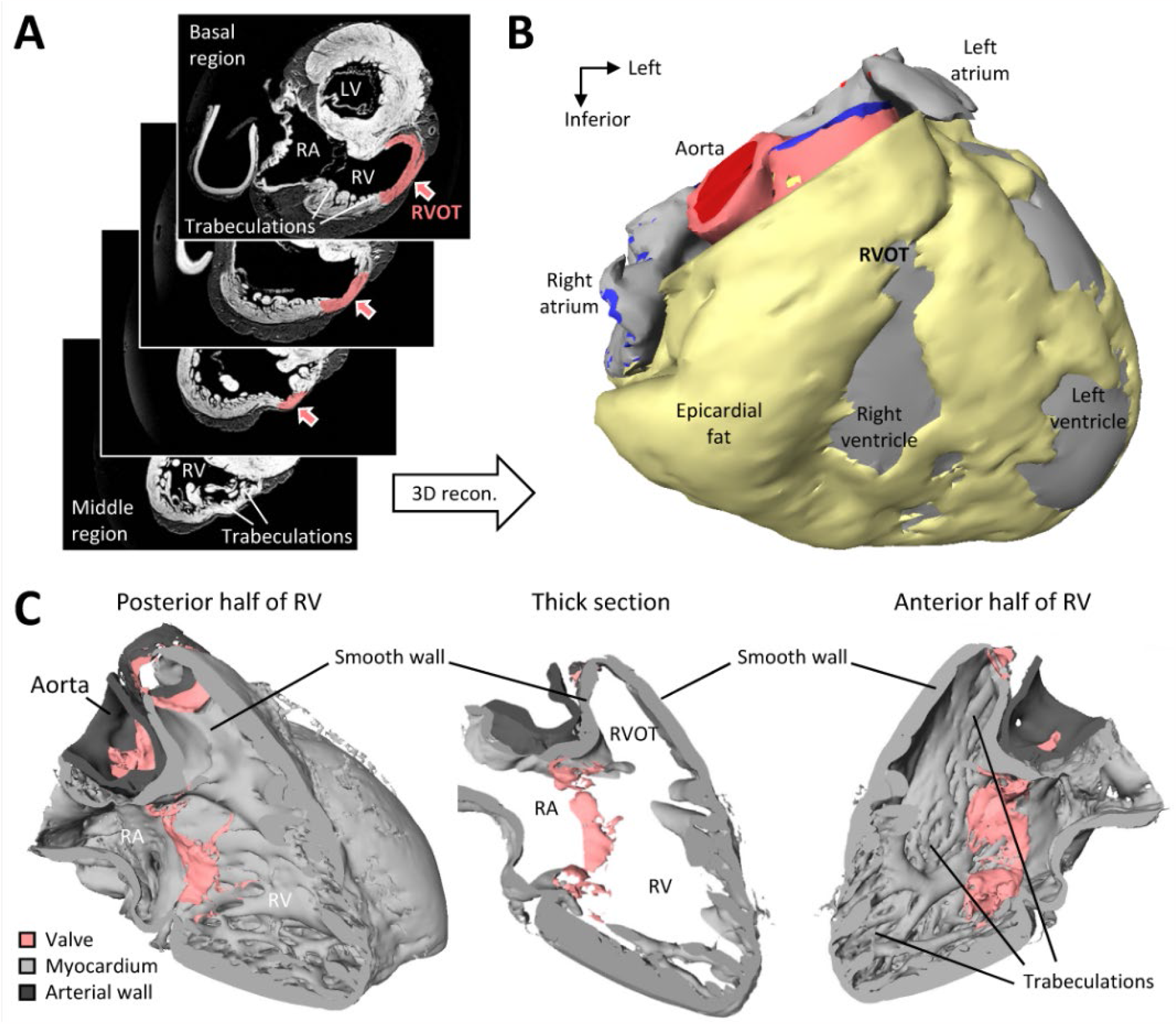
The outflow tract of the right ventricle. **A**. MRI-generated short-axis sections of a normal adult human heart, used to create a 3D model of the heart. The smooth, or a-trabecular wall, marked in red, characterizes the right ventricular outflow tract (RVOT). **B**. Anterior view of the 3D model (the interactive version is in Supplemental Materials). The RVOT myocardium is partially covered by epicardial fat in the reconstruction. **C**. Interior view of the right ventricle (RV) showing that the RVOT constitutes a substantial part of the right ventricle. LV, left ventricle; RA, right atrium; RV, right ventricle.

**Figure 2.**
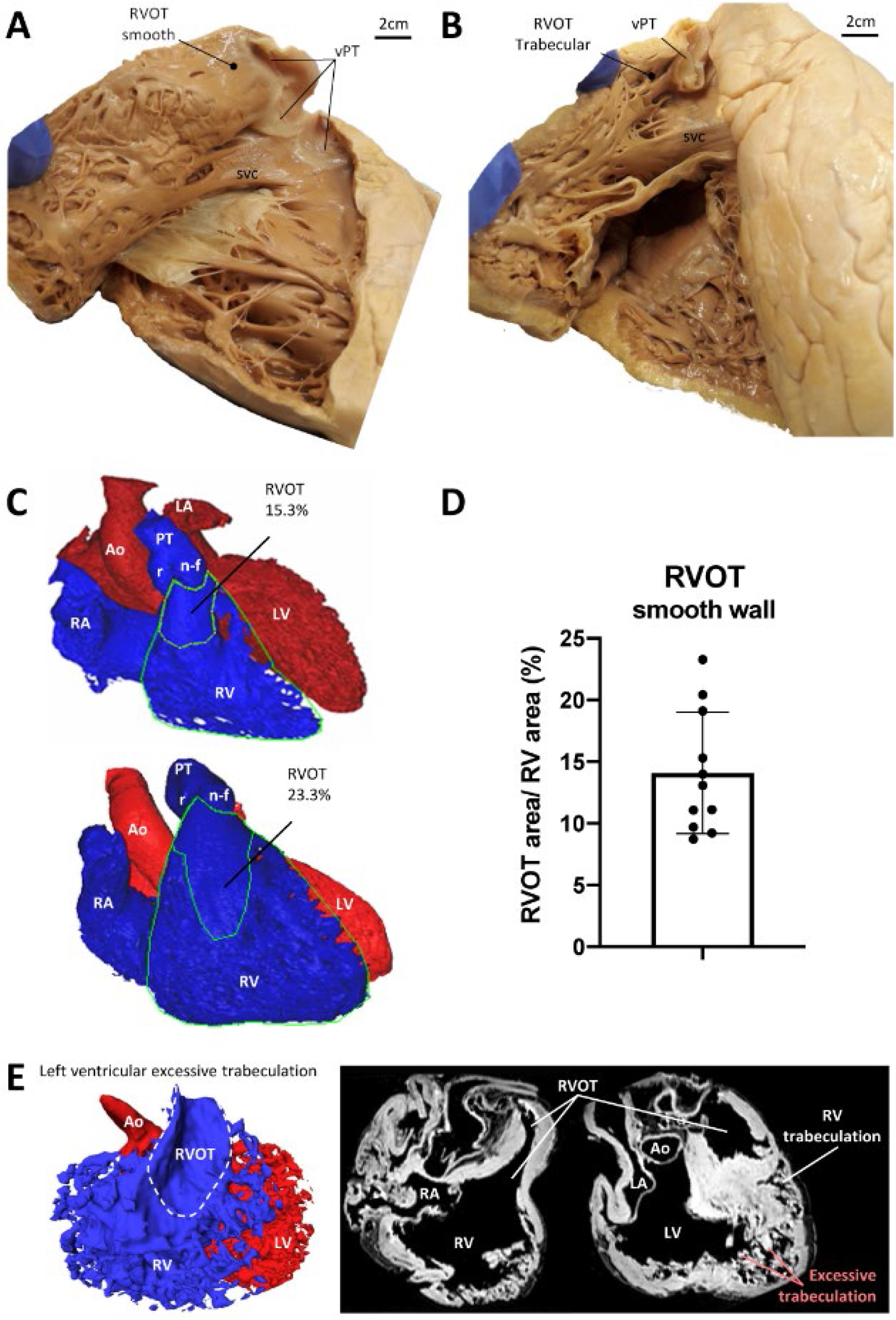
Morphological variation of the right ventricular outflow tract. **A**. Adult human heart with normal RVOT. **B**. Adult human heart with an abnormally high number of trabeculations in the RVOT. **C**. Anterior view of two virtual lumen casts in which the RVOT was measured to be 15.3% and 23.3% of the total right ventricular area (green lines indicate the measured areas). **D**. Graphical summary of all measurements of the relative size of the RVOT, showing the considerable variation between hearts (N=11). **E**. A case of left ventricular excessive trabeculation, showing a large RVOT. Ao, aorta; LA, left atrium; LV, left ventricle; PT, pulmonary trunk; RA, right atrium; RV, right ventricle; RVOT, right ventricular outflow tract; svc, supraventricular crest; vPT, pulmonary valve.

### Histological characterization shows prominent transmural variations in the RVOT

Using photographs of automatic serial sectioned cryo-preserved human RVOT myocardium we visualized a subendocardial longitudinal layer of myocardium and subepicadial oblique myocardial bundles (Figure 3A). This was corroborated using Masson’s trichrome staining of sections from six other hearts and it aligns with previous descriptions (17) (Figure 3B). Towards the ventricular apex, the oblique layer becomes continuous with the compact layer of the right ventricular free wall, whereas the longitudinal layer mostly gives way to trabeculations. The right ventricular wall has been measured to be 4 mm thick by echocardiography (18) and we found a similar thickness on histology of the right ventricular free wall and RVOT when the combined thickness of the compact and trabecular layers were considered (4.5±0.7 mm vs 4.2±.6 p =ns). The compact layers alone were not different in thickness between the right ventricular free wall and RVOT (2.8±0.5 mm vs 2.6±0.6 p =ns). The RVOT compact myocardium extends superiorly beyond the base of the pulmonary arterial valve (Figure 3B). This myocardium makes the so-called turret which was approximately 4 mm in both base width and height (Figure 3B). The attachment of the myocardium to the pulmonary arterial wall was slightly longer, approximately 5 mm (Figure 3B). In 5 of the 6 hearts studied, the RVOT was attached to the pulmonary arterial wall by myocardium following a longitudinal orientation. In the sixth heart, the attachment was 4.5 mm long, with the longitudinal myocardium attaching at 3.6 mm (80% of the length) and the transverse myocardium attaching at 0.9 mm (20% of the length).

**Figure 3.**
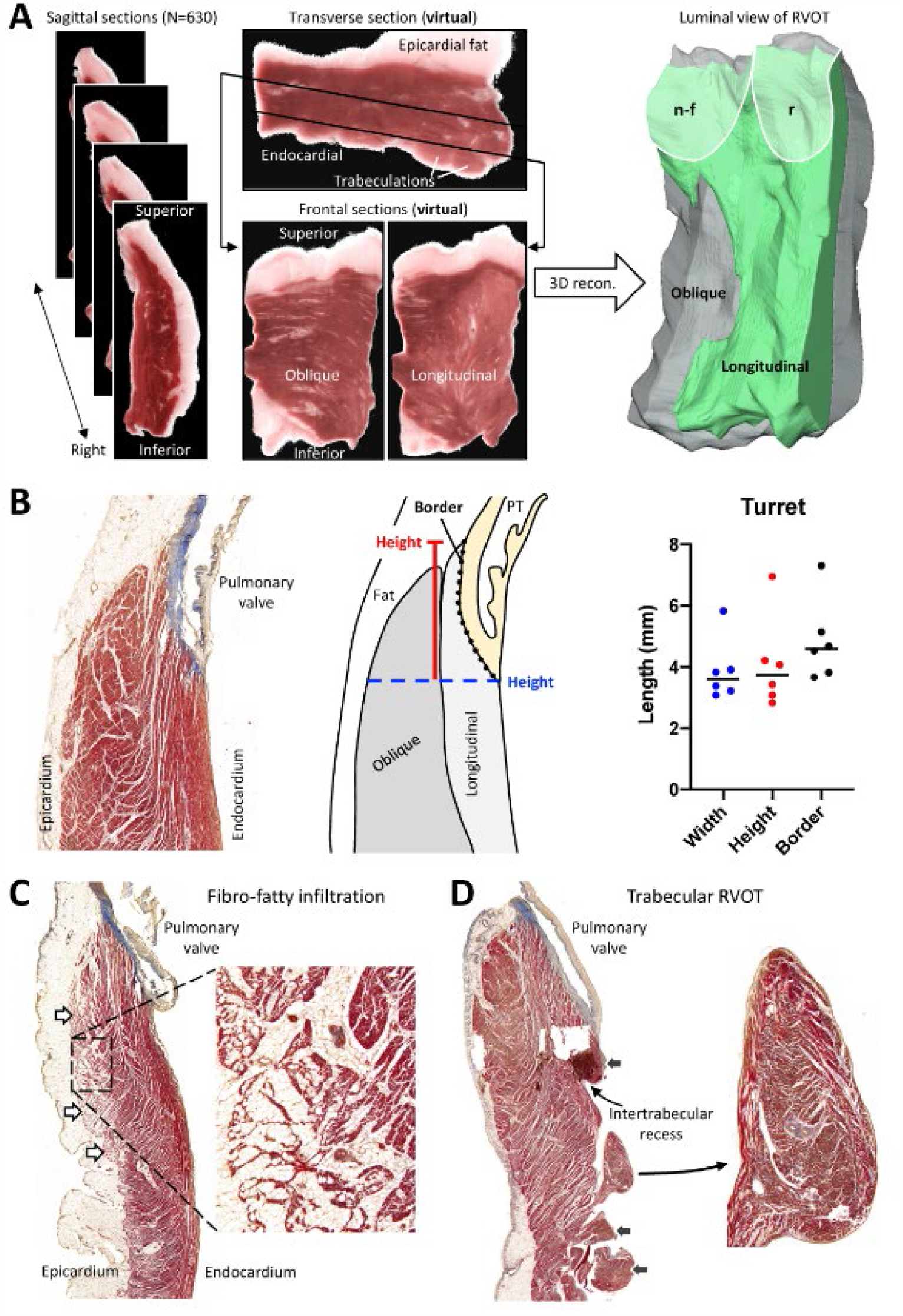
Histology of the outflow tract of the right ventricle. **A**. On the left, serial sections of bright field photographs of the human RVOT. The middle section illustrates the direction of myocardium at various locations. On the right, 3D reconstruction of the RVOT indicating the change in myocardial orientation from the luminal side to the epicardium. **B**. Measurements of the RVOT turret. The bar graph shows the morphometrics of the turret, N=6. **C**. A RVOT with much dispersion of the myocardium due to pronounced fibro-fatty infiltration from the epicardial side (white arrows). **D**. RVOT in which trabeculations (black arrows) extend to the base of the pulmonary arterial valve and the compact wall can be thin. n-f, non-facing leaflet of the pulmonary valve; PT, pulmonary trunk; r, right-facing leaflet of the pulmonary valve.

Some of the histological findings revealed that the RVOT myocardial wall contained transmural infiltrations of adipose tissue. In the example shown in Figure 3C, this fatty infiltration is pronounced in the epicardial half of the wall. While the fatty infiltration was most pronounced around coronary vasculature, it was found throughout the subepicardial RVOT myocardium. To assess myocardial density, histology images were converted to grayscale allowing quantification of myocardial density throughout the RVOT wall (N=6). No significant difference in myocardial density between both halves was found between epicardial and endocardial halves. Figure 3D shows an example of the RVOT where trabeculations extend to the base of the pulmonary arterial valve.

### Genes associated with trabecular myocardium are enriched in the subendocardium of the RVOT

To independently assess our anatomical observations of trabeculations on the luminal side of the RVOT on a molecular level, we collected fresh tissue samples from the RVOT subendocardium and subepicardium of 4 human hearts and performed RNA sequencing (Figure 4A). By differential gene expression analysis, we found 88 genes enriched in the subepicardium and 82 genes were enriched in the subendocardium (P adjusted < 0.05; Figure 4A). Subsequent gene ontology (GO) term enrichment analysis (16) yielded a subendocardial enrichment for “cardiac conduction”(Figure 4B). In the subepicardium, we found genes related to ‘developmental biology’ and ‘cellular response to stress’ to be enriched. The RVOT subepicardium showed higher abundance of biological processes associated with mitochondrial development and energy metabolism. In addition, several genes associated with trabecular development, e.g. *NPPA, IRX1, IRX3* and *IRX5*, were more abundantly expressed in the subendocardium of the RVOT (Figure 4C).

**Figure 4.**
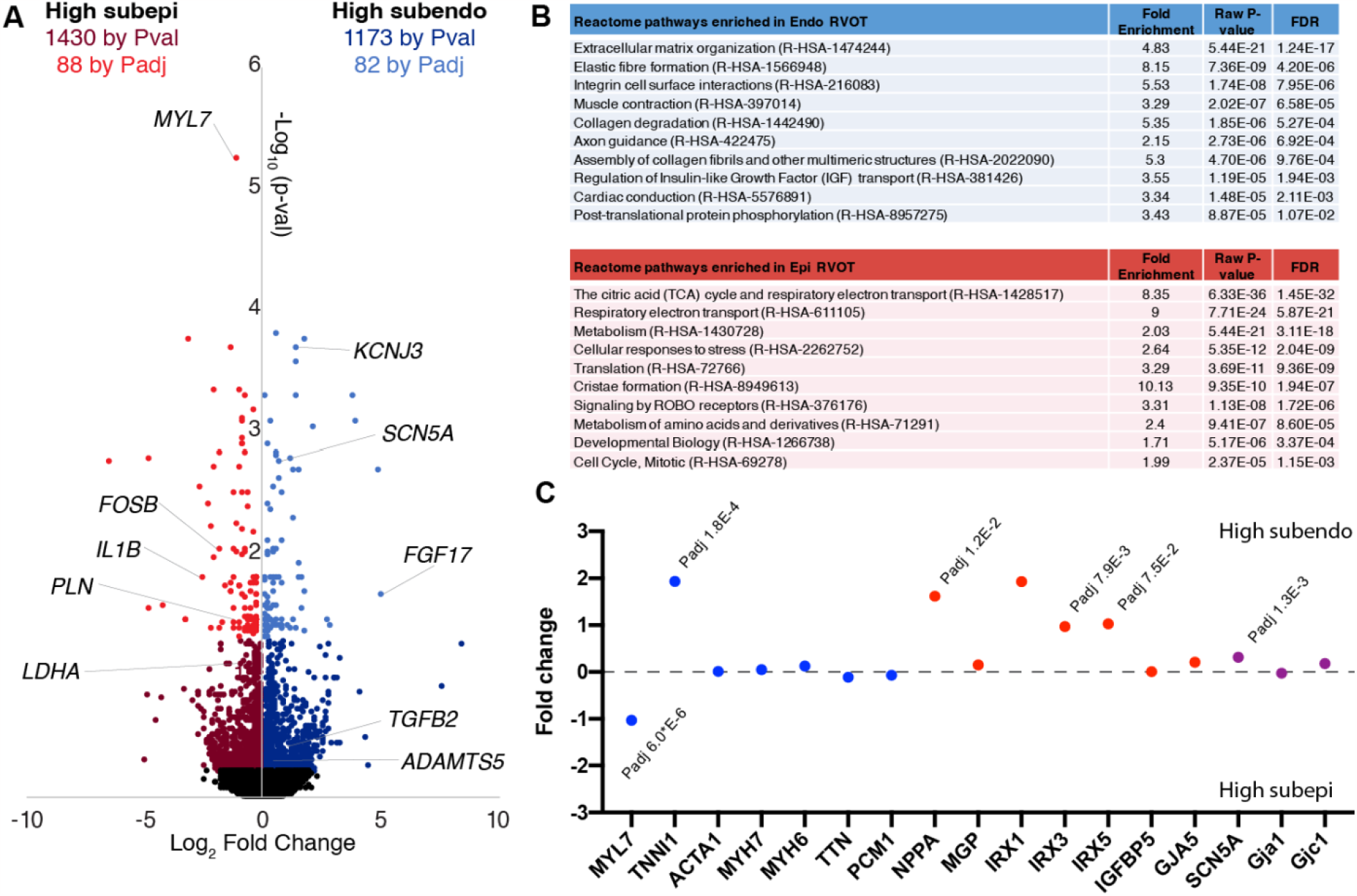
Gene expression analysis of the transmural wall of the right ventricular outflow tract. **A**. Scatter plot showing for each gene the statistical significance on the y axis and the magnitude of change on the x axis. **B**. Table showing the gene ontology analysis for gene enriched in either the subendocardium (blue) or the subepicardium (red). **C**. Graph showing selected genes involved in sarcomeric complex buildup (blue), development of trabecular myocardium (red), and in conduction (purple).

### Trabeculation of the luminal side of the RVOT predisposes to conduction delay

To investigate the effect of trabecular myocardium on impulse propagation in the RVOT, we recorded local bipolar electrograms in RVOT wedge preparations from two human donor hearts that showed pronounced trabeculations in the RVOT (Figure 5A). In both preparations fractionated electrograms could be recorded from the regions of the RVOT that harboured trabeculations. Figure 5B shows data from one heart illustrating the decrease in deflection amplitude and the decrease in the number of deflections when we stimulated with a shorter cycle interval. Simultaneously, optical mapping showed that the increase in fractionation coincided with an area of late activation close to the recording electrode which was placed in a region of the RVOT with trabeculations (Figure 5C-D). This activation delay was observed in both preparations. Altogether these electrophysiological findings provide evidence that the anatomical presence of the trabecular myocardium can form a substrate for hampered impulse propagation, which is potentially arrhythmogenic.

**Figure 5.**
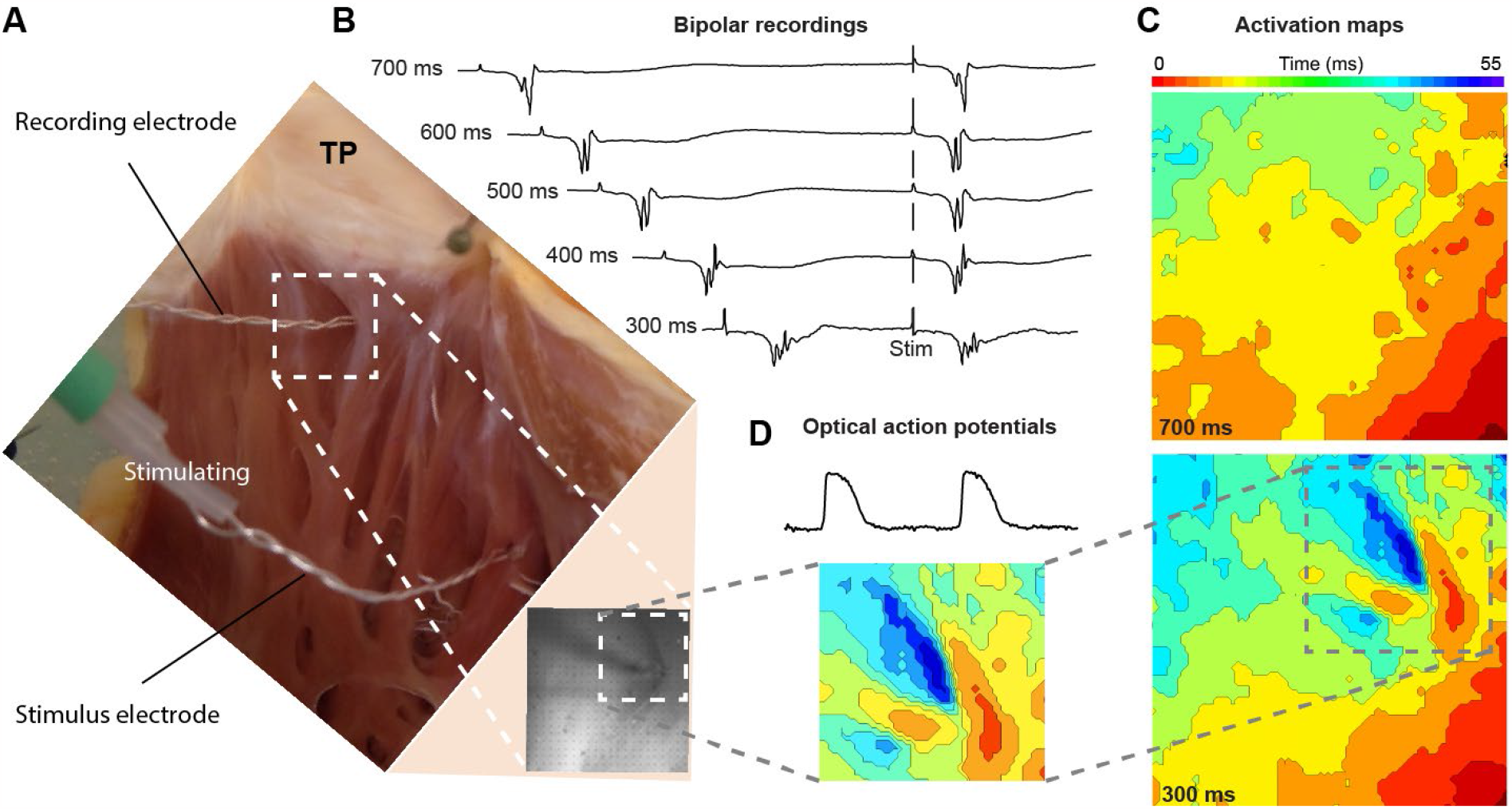
Trabecular myocardium in the RVOT may cause activation delay. **A**. A photograph showing the endocardial side of the RVOT wedge preparation during the experiment. White inset shows field of view during optical mapping **B**. Traces show bipolar electrograms at different pacing intervals recorded near trabeculations. **C**. Activation maps based on optical action potentials recorded at 700 ms (upper map) and 300 ms (lower map) intervals. The grey inset indicates the location of the trabecular meshwork near the recording electrode. **D**. Top: optical action potentials and bottom: activation map showing the activation delay over the trabecular meshwork

## Discussion

The anatomy of the RVOT is known to vary between individuals (19), but the marked variation in RVOT trabeculation that we document here has not been recognized until now. Such variation is likely to be related to the magnitude of key developmental processes and has potential implications for arrhythmia risk and ablation treatment.

The non-smooth-walled luminal portion of the RVOT has trabeculations of variable size and number. Ventricular trabeculations develop in the embryo under the control of complex signaling involving the endocardium (20). They are also the substrate from which the Purkinje cells of the ventricular conduction system develop (21). In the cow, and probably in other ungulates as well, the Purkinje network extends to the RVOT and almost reaches the pulmonary valve (22). In human, the Purkinje network is less penetrant than in ungulates, but the presence of Purkinje cells in the RVOT have been claimed based on histology (23). Our RNA sequencing results are consistent with the presence of Purkinje cells on the endocardial side of the RVOT. Consequently, this raises the possibility that a highly trabeculated RVOT may also contain more Purkinje cells than a smooth-walled RVOT. This relationship should not be overstated, as a state of excessive trabeculation in humans does not overtly affect the Purkinje system and ventricular arrhythmias do not preferentially originate from the excessive trabeculations (24). Nevertheless, in so far the presence of Purkinje cells influences the risk of arrhythmogenesis, the extent of trabeculation of the RVOT may be clinically relevant.

The presence of trabeculation in the RVOT appears to affect the thickness of its compact wall. Put simply, the RVOT wall is approximately 4 mm thick, and in a setting of trabeculation, a smaller proportion of the 4 mm is made up of the compact layer. This is similar to the setting of excessive trabeculation, where a thicker trabecular layer is associated with a thinner compact layer (25). A thinner compact layer provides a thinner barrier between the right ventricular cavity and the pericardial space. The depth of the compact wall that can be surgically ablated in the treatment of ectopic pacing or reentrant tachycardia will be less in a trabeculated RVOT. Many trabeculations are much smaller than what can be visualized with typical clinical echocardiography and MRI (7, 12) and contrast-enhanced CT may give a more accurate reading of the depth of tissue that can be ablated.

Our histological findings confirm that the anterior RVOT compact wall is made up of two layers of myocardium aligned in a circular to oblique fashion subepicardially and aligned in a longitudinally fashion subendocardially (19). Our results also show that this myocardium can be quite dispersed by fibro-fatty infiltration. As a result, the myocardium is divided into discrete bundles, giving rise to discontinuous conduction (26). Whether fibro-fatty infiltration is the primary source of myocardial discontinuities in the RVOT or merely extends pre-existing discontinuities is not clear. As we observed mainly fatty infiltration rather than fibrosis, this is most likely a common finding in aged hearts rather than arrhythmogenic right ventricular dysplasia (27). Nevertheless, the structural integrity and conduction properties of the RVOT may be compromised, resulting in an altered propensity for arrhythmogenesis (28).

The RVOT forms in the first trimester, together with most cardiac structures (29). It is derived from the slowly conducting embryonic outflow tract (OFT), which in turn is derived from cardiac precursors in the second heart field (30). This process is regulated by the T-box transcription factor 1 (TBX1) (31). It is currently unclear whether altered expression levels of TBX1 affect the size of the right ventricular myocardium derived from the embryonic OFT, i.e. the RVOT and parts of the RV. We speculate that the size of the smooth wall portion of the right ventricle is related to the extent to which the embryonic OFT is incorporated into the right ventricular wall. This ventricularization of the OFT can be considered complete when trabeculations begin to form in what was previously smooth-walled outflow tract myocardium. In this sense, the variation in size and length of the smooth-walled part of the RVOT is plausibly related to the extent of embryonic morphogenetic processes (9). Interestingly, the extent of RVOT trabeculation in the adult heart, is likely without correlation to the extent of left ventricular trabeculation. Taken together, while variation in trabeculations of the RVOT can be readily observed, how and when this variation arises is mostly obscure.

## Conclusion

Trabeculations are present in the human RVOT with varying degrees among individuals. As a result, the thickness of the compact wall in the RVOT is affected, which in turn limits the depth of tissue accessible for clinical interventions and influences electrical propagation, setting the stage for arrhythmias.

## Conflict of Interest

The authors have no conflicts of interest to declare.

## Acknowledgments

We would like to thank Roelof-Jan Oostra for providing access to human hearts of the Becker collection (AMC, Amsterdam UMC), Manuela Frimpong for assistance with histology.

## Data availability

The data are available at https://github.com/bjboukens/Data-Jensenetal

## Notes

https://github.com/bjboukens/Data-Jensenetal

### Competing Interest Statement

The authors have declared no competing interest.

