## Supplementary Figure 1 for "The smooth-walled human RVOT contains trabeculations that cause conduction delay"

### Information on the use of this interactive 3D-PDF

All labels
- Right lumen
- Right av valve
- RV compact layer
- RV trabecular layer
- RV outflow tract
- Pulmonary trunk
- Pulmonary valve
- Left lumen
- Left av valve
- Aorta
- Other tissues
- Epicardial fat

Selected structure:

Left lumen

**Selection of structures**

The top left panel contains buttons to show or hide (groups of) structures, or to make them transparent.

**button example:** show / transparent / hide — Right av valve

After a single click on a 3D structure, the structure will be highlighted and the name of the structure will appear below "Selected structure". With the buttons next to the structure name, the appearance of this structure can be changed. Clicking next to the 3D object will deselect the structure.
For more advanced selection options, right-click on the 3D model and choose: “Show Model Tree”.

**The intended use and characteristics of this model**

This 3D reconstruction is intended to facilitate the understanding of the morphology and position of the human right ventricular outflow tract (RVOT). In this particular heart, a fairly typical RVOT is found. The basis for the reconstruction is an ex vivo MRI scan with a spatial resolution of 0.4 mm in each plane. Such resolution allows for the reasonably accurate reconstruction of larger structures, such as the RVOT. Thin structures such as the atrioventricular valves, in contrast, are on the border of what can be faithfully reconstructed. The smallest trabeculations are approximately as thin as valve leaflets, and it should be presumed that the smallest trabeculations are not reconstructed in this model. There were no major lesions in this heart. The state of contraction has not been controlled and while the right ventricle is approximately diastolic, the left ventricle is much contracted. Much of the atria is included, but the superior part is missing.

#### Technical Notes

View this PDF file in a recent version of Adobe Acrobat Reader: https://get.adobe.com/reader/

3D interaction is only possible on MS Windows or Mac OS. Javascript and playing of 3D content must be enabled.

*Edit, Preferences* to ensure the following:

1) In *JavaScript*
   - enable *Enable Acrobat JavaScript*
2) In *Multimedia &3D*
   - enable *Enable playing of 3D content*
   - disable *Show 3D Orientation Axis*
   - *Optimization Scheme for Low Framerate: None*

Amsterdam UMC
University Medical Centers

Department of Medical Biology

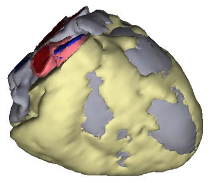

Frontal view

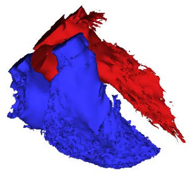

Lumen cast

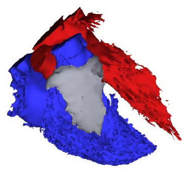

RV outflow tract

### Reconstructed 3D model of a human heart

All labels
- Right lumen
- Right av valve
- RV compact layer
- RV trabecular layer
- RV outflow tract
- Pulmonary trunk
- Pulmonary valve
- Left lumen
- Left av valve
- Aorta
- Other tissues
- Epicardial fat

Selected structure:

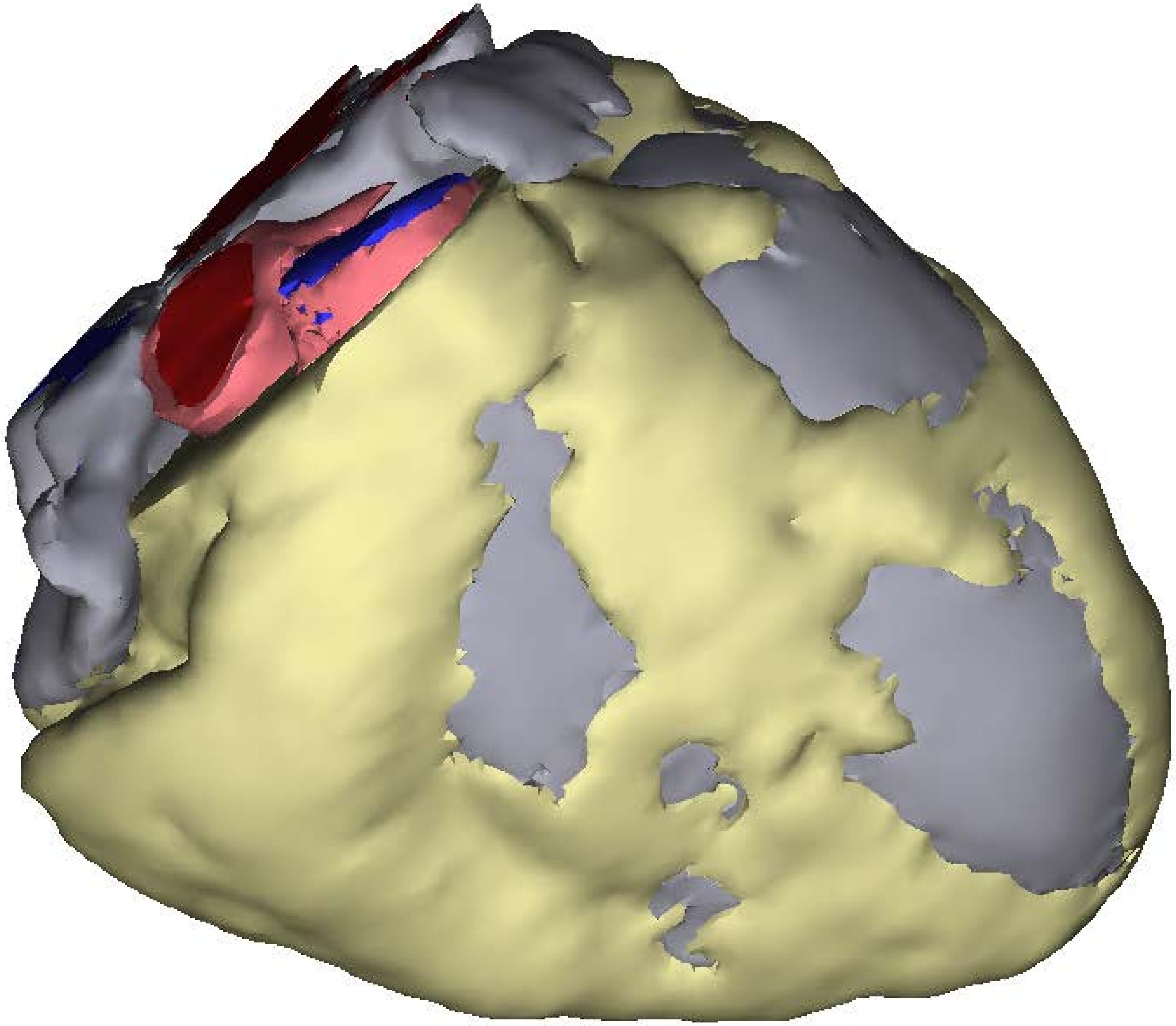

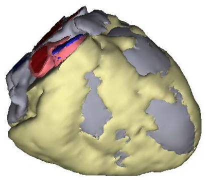

Frontal view

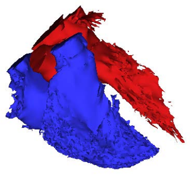

Lumen cast

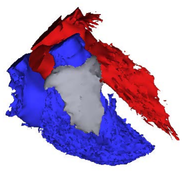

RV outflow tract

Amsterdam UMC
University Medical Centers

Department of Medical Biology
