## Supplementary tables for "The smooth-walled human RVOT contains trabeculations that cause conduction delay"

Supplementary Table 1. Data on heart volume renderings from Atlas of Human Cardiac Anatomy. F, female; M, male.

| Specimen | Sex | Age |
| --- | --- | --- |
| Heart0110 | F | 61 |
| Heart0141 | F | 24 |
| Heart0148 | F | 73 |
| Heart0188 | F | 61 |
| Heart0215 | F | 57 |
| Heart0096 | M | 54 |
| Heart0111 | M | 25 |
| Heart0131 | M | 51 |
| Heart0145 | M | 53 |
| Heart0094 | ? | ? |
| Heart0102 | ? | ? |

Supplementary Table 2. Data of patients of whom hearts were used for quantitative morphology. F, female; M, male.

| Specimen | Sex | Age | Cause of death |
| --- | --- | --- | --- |
| #1 | M | 33 | Liver angiosarcoma, complicated by acute hemorrhage |
| #2 | M | 51 | Pancreatic adenocarcinoma with cerebral metastasis |
| #3 | F | 60 | Pulmonary adenocarcinoma with widespread metastasis |
| #4 | M | 67 | Cerebral tumor |
| #5 | F | 77 | Cerebrovascular accident |
| #6 | F | 92 | Intestinal hernia incarcerate, followed by sepsis |
